# Methodological priorities in assessing wild edible plant knowledge and use – a case study among the Baka in Cameroon

**DOI:** 10.1101/2020.05.20.106427

**Authors:** Sandrine Gallois, Thomas Heger, Amanda G. Henry, Tinde van Andel

## Abstract

Freelisting and dietary recalls are frequently used methods in ethnobotany to assess wild edible plant (WEP) knowledge and use. Though these *ex-situ* interviewing methods are practical to perform and may yield large datasets in a short time, they are known to be limited by the informant’s memory and cognitive bias. Alternatively, the much more laborious walk-in-the-woods method may be used, in which informants point out edible plants *in-situ*. Few studies, however, examine quantitatively how these different methods influence results. In this study, we assessed how these methods capture the diversity of wild edible plant knowledge and use among the Baka, a group of forager-horticulturalists from southeastern Cameroon. We show that within a single population, and when data on consumption frequency are collected simultaneously, the walk-in-the-woods method results in more detailed information of WEP knowledge and use than do freelisting or dietary recalls. Our *in-situ* method yielded 91 species of WEP, much more than the *ex-situ* methods of freelisting (34 spp.) and dietary recalls (12 spp.). Our results imply that previous studies based only on *ex situ* surveys may have underestimated the importance of WEP for local communities. We propose that future studies on WEP knowledge and use frequency should rely on mixed methods, taking an *in-situ* method as the starting point of their approach.

## Introduction

The value of local ecological knowledge in informing conservation and environmental management is well established (Chazdon et al., 2009; Pandey & Tripathi, 2017; Pardo-de-Santayana & Macia, 2015). Local knowledge on useful plants may be especially valuable in this regard (Cummings & Read, 2016). For example, in an ethnobotanical study in Rio Formoso, Northeastern Brazil, Da Cunha and De Albuquerque (2006) found that the main product harvested from over half of the useful plant species was wood, indicating the need for conservation initiatives to provide an alternative for this source of fuel and construction material. Additionally, ecological knowledge is also crucial for local people, especially those who intimately depend on their surrounding natural resources for their subsistence and who have developed through generations a substantial expertise on the use of wild plants and animals for food, shelter and medicine (Reyes-García, 2015). However, the access and availability of natural resources central for dietary diversity and food security of these societies are challenged by natural resources exploitation, such as mining and selective logging, commercial harvesting and hunting, especially in areas where the biodiversity is high such as tropical rainforests (Baudron et al., 2019; Wasseige et al., 2012). These pressures on local resources has led to a decrease in local people access to the important wild plants and game that contribute to their diet and medicine (Rist et al., 2012). Therefore, considering the different potential conflict of use among wild resources, accurate assessments of the use of wild plants are necessary in order to evaluate the effects of overall global changes affecting both biodiversity and local livelihoods.

Local knowledge and use of plants are assessed through different ethnobotanical methods, of which the interview is the most widely used. Different interview methods are deployed based on the research question addressed and vary considerably between studies (Thomas et al., 2007). Freelisting is a frequently-used method, in which informants are asked to list all items they know within a given category (Martin, 2010). This technique reveals cultural salience and variations in individuals’ topical knowledge (Quinlan, 2005), and results in a shortlist of highly valued plants (Ghorbani et al., 2012; Mengistu & Hager, 2008). As freelisting allows the collection of data from a large number of informants in a limited amount of time (De Sousa et al., 2016), this method is frequently used as a starting point for studying traditional plant knowledge. Plants listed during the interviews are collected and identified afterwards. The resulting dataset is then used to draw conclusions about plant knowledge of a certain group of people and/or the potential contribution of wild plants to their diet (Fongnzossie et al., 2020; Mengistu & Hager, 2009; Termote et al., 2011). Organizing field trips to collect herbarium specimens of species mentioned during freelisting exercises is often (inaccurately) called the ‘walk-in-the-wood method’ (Lulekal et al., 2013; Termote et al., 2011). However, this technique, first coined by Phillips and Gentry (1993), implies that participants are encouraged to actively lead field trips and point out all useful plants they know and/or use (Thomas et al., 2007), instead of only searching for specimens that appear on the list of local names derived from interviews.

Data elicited from freelisting appear to be specific to the context in which they were collected (e.g., in the village), creating an unintended but significant bias in this type of ethnobotanical research (De Sousa et al., 2016; Martin, 2010; Paniagua Zambrana et al., 2018). Gathering the data *ex-situ* (away from the ecological context in which people collect their plants) may result in lists of only the most salient plant species. Furthermore, the success of freelisting depends on the informants’ correct understanding of the category or cultural domain (e.g., wild food plants) under discussion (Da Cunha & De Albuquerque, 2006; Quinlan, 2005; Quiroz et al., 2016; Gallois et al. 2020).

In societies that undergo rapid socio-economic changes, people become more integrated into the market economy, change their lifestyle and adopt cultivated or processed substitutes for wild plants in their diet (Kuhnlein, 2009). This creates a gap between people’s ethnobotanical knowledge and their actual use of plants (De Albuquerque, 2006; Reyes-García et al., 2005). A discrepancy between the number of useful species known and those actually used indicates that elders who still know how plants were used in the past do not practice this any longer, and infrequently transfer their skills to the next generation (Reyes-García et al., 2005). Freelisting exercises often focus on peoples knowledge (Reyes-García et al., 2005) while recall surveys, developed by social anthropologists for understanding time allocation (Gross, 1984), lead informants to enumerate what they have done during a specific period of time. Recently, recall surveys were introduced in ethnobotanical approach to assess local uses of plants. For instance, dietary recall surveys have been developed to estimate the proportion of different food items in people’s diet (e.g., Munger et al. (1992); Friant et al. (2019); Reyes-García et al. (2019)), while income recall survey have been used for assessing the contribution of the sale of different forest products in local livelihood (see for instance Levang et al., (2015)). Although many studies reported a high diversity of wild edible plant species worldwide (Bharucha & Pretty, 2010; Delang, 2006), research relying on dietary recalls has also resulted in surprisingly low numbers of wild species actually being consumed (do Nascimento et al., 2013; Ogle, 2001).

In the highly biodiverse context of the Central African Congo Basin, a wide variety of wild edible plant species has been reported by Bantu-speaking farmers (Ingram & Schure, 2010; Termote et al., 2011; van Dijk, 1999) but especially among hunter-gatherers that infrequently practice agriculture (Bahuchet, 1992; Dounias, 1993; Ingram & Schure, 2010; Terashima & Ichikawa, 2003; Yasuoka, 2012). Dietary recalls carried out in the Democratic Republic of the Congo, however, showed that wild plants did not contribute substantially to rural and urban women diets (Termote et al., 2012). Likewise, dietary recalls held among the Baka people in Cameroon resulted in only 15 wild edible species being reported (Gallois et al., 2020), which is in stark contrast to the extensive wild plant knowledge reported earlier by Bahuchet (1992) and Dounias (1993) for the same ethnic group. Like freelisting, dietary recalls are limited by the subject’s memory (Grandjean, 2012) and may therefore underreport plant use. In this study, we explore how different ethnobotanical methods capture the diversity of wild edible plant knowledge and use among a community of Baka forager-horticulturalists in southeastern Cameroon. We aimed to answer the following questions:

1. Which wild edible plant species (WEP) are reported by the Baka during freelisting, dietary recalls, income recalls, and walk-in-the-woods methods?
2. How do the results differ between these methods?
3. What are the general characteristics of the WEP known and consumed by the Baka?
4. How do conclusions based on the results obtained by the four methods differ in terms of the potential conflicts in use among local consumption, logging, and trade?

We hypothesized that walk-in-the-woods would result in a larger number of plant species than the other three methods, but that all four methods would identify the species most frequently consumed by our informants. We also predicted that the list of plants given through freelisting, and dietary and income recalls would underestimate the potential conflicts in use of edible plants.

## Methods

### Study site

Data were collected around the villages of Le Bosquet (3°07’38’’N13°52’57’’E) and Kungu (3°02’40“N 14°06’57”E), located in the Haut Nyong division, southeastern Cameroon. The communities are located at least eight hours by car from the capital Yaoundé, of which four hours are on unpaved logging roads. The accessibility of this area highly depends on the weather, as the road quickly deteriorates during the rainy season. The area is covered by a mixture of evergreen and moist semi-deciduous forest within altitudinal ranges of 300–600 m. (Letouzey, 1985). In populated areas, the forest cover is largely removed in favor of settlements, cocoa plantations, logging activities and small-scale agriculture. This creates a mosaic of dense primary forest, selectively logged primary forest, secondary forest and agricultural fields, interspersed with trails. The climate of the region is tropical humid, with a major rainy season between late-August and late-November and a major dry season between late-November and mid-March. The annual precipitation reaches about 1500 mm and the average temperature is 25°C (Leclerc, 2012).

The area is populated by two main ethnic groups: the Nzimé, Bantu-speaking farmers, and the Baka, Ubangian-speaking forager-horticulturalists. Until roughly 50 years ago, the Baka were nomadic foragers, relying on hunting, fishing, gathering, and the exchange of non-timber forest products against agricultural crops with their farming neighbors. Since the 1960s, the Baka have been facing several changes in their livelihood. Due to a government program of sedentarization (Leclerc, 2012), they have progressively left their forest camps and settled in villages along the logging roads. Nowadays, their livelihood is mostly based on the combination of foraging activities, agricultural work in their own fields and wage labor for the Nzimé or for logging companies (Gallois et al., 2020).

### Data collection

We used a combination of four different datasets, obtained from freelisting, dietary recalls, income recalls, and ethnobotanical field surveys. Data collection took place in both villages in three different fieldwork periods: February-March 2018 (major dry season), October-November 2018 (major rainy season), and April-May 2019 (minor dry season) to cover variations in wild fruit availability. The freelisting data were gathered during the first fieldwork period, income recall data during the two first fieldwork periods, and dietary recall data during all three fieldwork periods. The walk-in-the-wood surveys were carried out during the last fieldwork period. Before data collection, Free Prior and Informed Consent was obtained from all participants. This study adheres to the Code of Ethics of the International Society of Ethnobiology (2006), received approval from the ethics committee of Leipzig University (196-16/ek), and the Ethical Committee from the Ministry of Health of Cameroon (n°2018/06/1049/CE/CNERSH/SP).

We conducted freelisting exercises among 55 Baka individuals of 18 years and older (24 men and 31women), during which we asked our interviewees to report all wild edible plants they knew (Gallois et al., 2020). We gathered data on the importance of wild plants in Baka diet by conducting a dietary recall protocol that was adapted from the FAO Guidelines for Assessing Dietary Diversity (Kennedy et al., 2011). Informants were asked to list all items they had consumed within the previous 24 hours, and to mention the origin of each food item (from the wild, from agricultural fields or bought at the market). A total of 143 dietary recall interviews were conducted among 83 informants (35 men and 48 women): 42 individuals were interviewed once, 22 twice and 11 three times. Finally, we also collected data on wild edible species that were traded as timber and as non-timber forest products. We conducted a 14-day recall survey on the income received through sale, asking our interviewees to list all the items they had sold during this time period. A total of 114 interviews were conducted over 34 individuals in le Bosquet and 39 in Kungu (in total 43 women and 30 men): 32 were interviewed once and 41 twice.

From the local names of wild edible plants mentioned during the different interview methods, we constructed a preliminary database of species consumed by the Baka, with tentative scientific names from literature on Central African wild food plants (e.g., (Bahuchet, 1992; Betti et al., 2013; Brisson, 2010; Dounias, 1993; Yasuoka, 2012) Finally, for our walk-in-the-woods trips, we asked the community to suggest several people of different ages and gender that were knowledgeable on wild edible plants and would agree to join us on our collection trips as informants. We worked with one to four informants on each collection day. In total, we employed 20 informants (10 women, 10 men, aged between 29 and 80 years). Nine informants had also participated in the previous *ex-situ* interviews (dietary and income recalls: 2; free listing: 2; all three methods: 5). During 14 collection days into the area surrounding Le Bosquet and Kungu we asked our informants to point out any edible plant they saw. We also searched for the species on our preliminary list of wild food plants. When a wild edible plant was encountered, herbarium material was collected using standard botanical methods (Martin, 2010). For most specimens collected, we asked our informants for 1) the local name in Baka (or French /Nzimé if known); 2) plant part(s) used; 3) preparation and application methods; 4) when they had last consumed the plant; 5) whether a part of the plant was sold;6) whether it was commercially logged. To analyze conflicts between commercial timber harvesting and the availability of wild food plants for the Baka, we documented the local names and we also counted the number of logged tree trunks along the forest trails and on logging trucks passing through the village.

Duplicates of voucher specimens were deposited at the National Herbarium of Cameroon (YA) and Naturalis Biodiversity Center (L). A third voucher was used in the study site to discuss local names and uses with Baka villagers. Plant identification took place at Naturalis, using Central African herbarium specimens and literature (e.g., Harris & Wortley (2018); Hawthorne & Carel Jongkind (2006); Hutchinson & Dalziel (1958); Royal Botanic Gardens Kew (1931-1973); MNHN (1963-2018)). This literature was also used to verify the vegetation types in which these WEP occurred naturally. For species that were difficult to identify, we consulted botanical experts at Naturalis and abroad. Scientific names were updated using the portal of Plants of the World Online^1^.

### Data analysis

In order to assess the differences in results coming from the different methods, we compared the total number of wild edible species encountered during the walk-in-the-woods, freelisting and dietary recalls. To assess whether the full potential of the methods had been utilized, species accumulation curves (Peroni et al., 2014) were produced for each of them by calculating the cumulative number of species that were reported after a certain amount of collection days (walk-in-the-woods method) and after interviewing a certain number of informants (freelisting and dietary recalls). Contrary to usual practice, data were not randomized before producing the curves, as several relevant features of the data would have been lost. As the income recalls were only used to assess commercialized WEP, a subset of all wild edible plants, we did not produce a species accumulation curve. To assess the general characteristics of wild species consumed by the Baka, information on life form, part used, habitat and commercial timber was categorized in a Microsoft Excel table, after which bar graphs were produced to show the distribution of these traits.

To analyze the actual use of WEP reported during the walk-in-the-wood trips, we first categorized the information of last consumption for each species according to Gallois et al. (2020) in the following categories: 1) today/yesterday; 2) within the week, 3) within the month; 4) within the year; 5) 1-2 years ago; 6) > 2 years ago; and 7) never. A bar chart was produced to visualize the ranking of the most recently consumed species and comparison of the results of walk-in-the-woods with the dietary recalls. We also compared the commercialized WEP reported during the income recall surveys to the plants said to be sold during the forest trips. Finally, we cross-referenced the species said to be cut for commercial timber and the CITES appendices^2^ and IUCN Red List^3^ to assess their current conservation status. Trade names of timber species were identified through vouchers specimens and the International Tropical Timber Organization’s website^4^.

## Results

### Capturing the diversity of edible plants: comparison between methods

The dietary recalls and freelisting resulted in 12 and 38 wild edible plant species respectively. Initially, 51 local names were identified through freelisting, but 13 of those were later excluded because they were either synonyms of Baka plant names that had already been mentioned (three names) or they referred to wild mushrooms (two names), types of honey (six names) or cultivated plants (two names). Two species that emerged from the dietary recalls (*Amaranthus dubius* Mart. ex Thell. and *Raphia* sp.) were not found through free listing. During the walk-in-the-woods method, we collected 94 wild edible plant specimens that corresponded to ca. 91 species, which included all species mentioned during the dietary recalls and freelisting methods. The exact number of wild edible species is unclear, as eight vouchers could only be identified at the genus level and for several West- and Central African *Dioscorea* species (wild yams), the taxonomic species delimitation is not clear (Magwé-Tindo et al., 2018).Moreover, the Baka recognize different forms within individual yam species and thus some local names refer to the same botanical taxon. For instance, in the case of *D. minutiflora*, the Baka distinguish three distinct types: “njàkàkà”, “bálOkO” and “kuku”, all with different leaf and tuber morphology. All local and scientific names of each wild edible species, used parts, preparation methods, consumption frequency and the method(s) through which they were recorded are listed in Supplementary Material.

Over the 83 individuals interviewed during dietary recalls, only 69 reported wild edible plants. The species accumulation curve for the dietary recall method approached the asymptote after interviewing 83 people (Figure 1). Between respondents 46 and 83, only three new species were mentioned, which suggests that interviewing more respondents would not have led to many more wild edible plant species being identified. Therefore, the dietary recall appeared to have captured most of the WEP diversity that was possible with this method.

**Figure 1.**
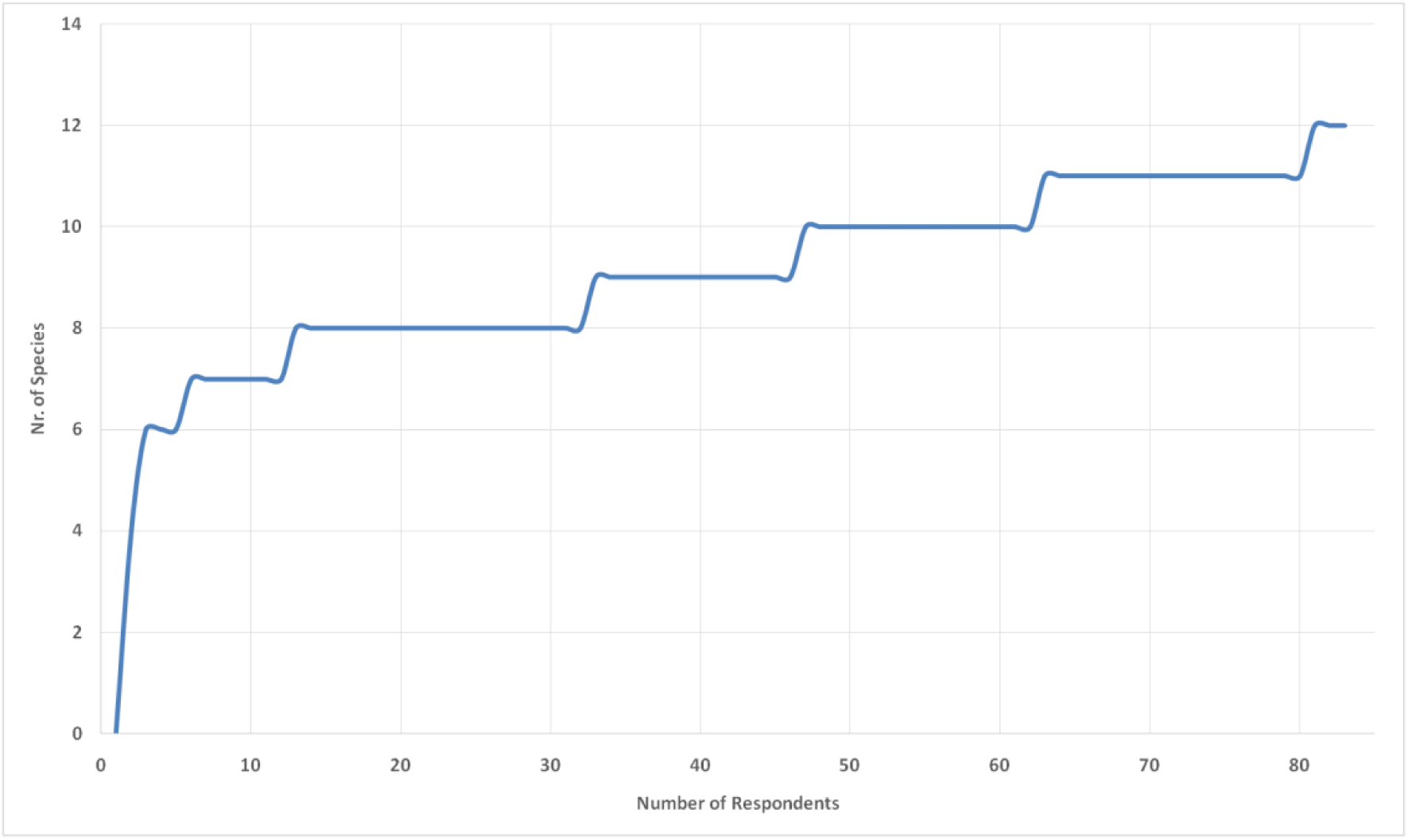
Species accumulation curve of wild edible plants mentioned during the 83 dietary recalls in Le Bosquet and Kungu, southeast Cameroon, 2018.

The species accumulation curve of the freelisting methods approached the asymptote after interviewing 55 individuals, with a total of 38 WEP species reported (Figure 2). This indicates this method also efficiently captured the requested information, at least within its limitations. Typically, 14 of the 55 respondents reported not knowing any wild edible plants, which resulted in several flat sections in the curve.

**Figure 2.**
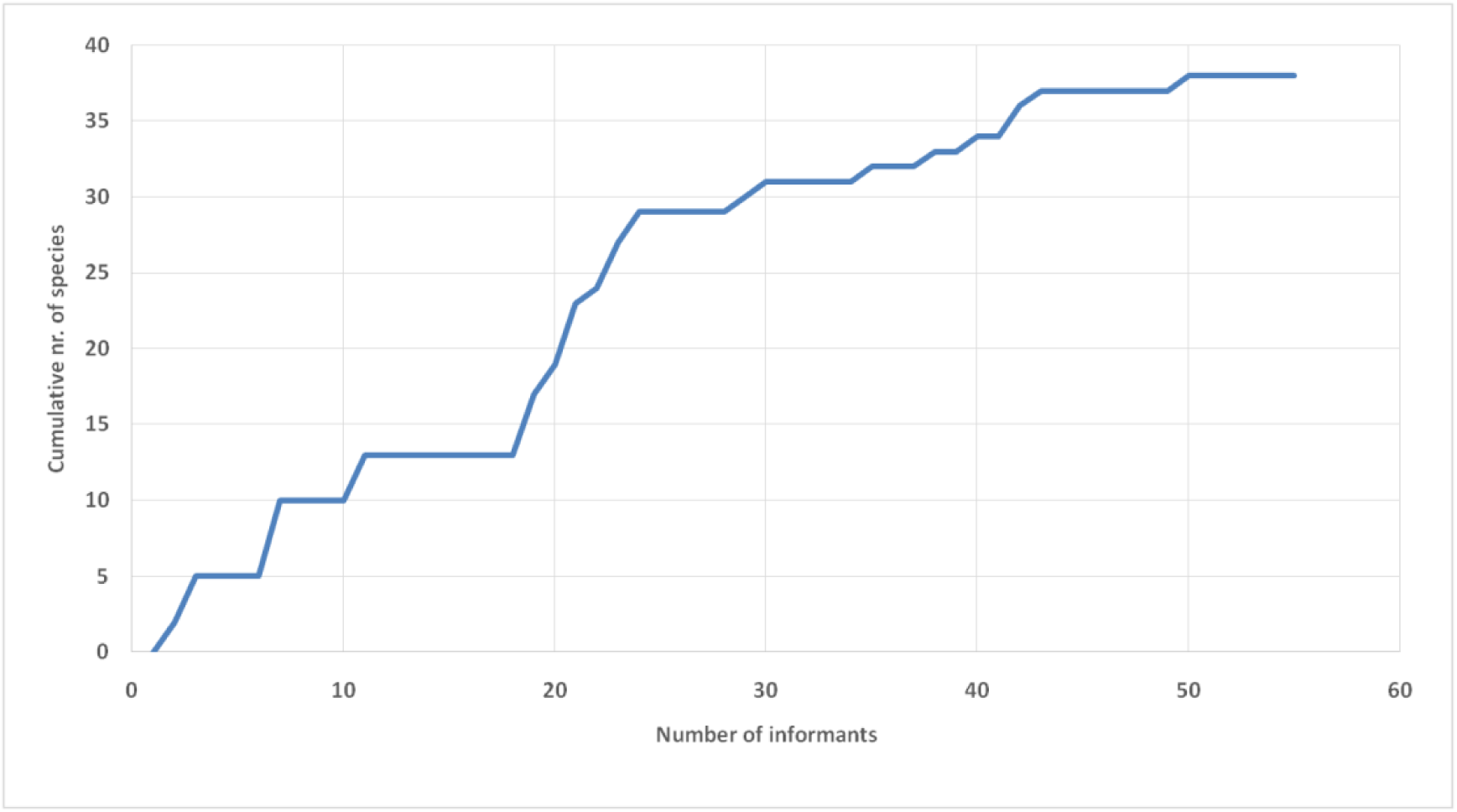
Species accumulation curve of wild edible plants mentioned during the 55 freelisting interviews in Le Bosquet and Kungu, southeast Cameroon, 2018.

The species accumulation curve for the walk-in-the-woods method flattened somewhat after 11 days, but not completely (Figure 3).This suggests that more WEP would have been recorded if fieldwork had continued. Our Baka informants indeed mentioned that there were additional rare species that could only be found after walking for hours in the forest. We know that at least four other species could have been found if we had more time to walk further into the forest. From their Baka names and the literature (Brisson 2010), we assume that these WEP were the African mammee apple (*Mammea africana* Sabine) with large edible fruits, a species of *Afzelia*, of which the red arils around the seeds are eaten, a species of *Raphia* palm tree of which the sap is fermented into palm wine, and the African walnut tree (*Coula edulis* Baill.) that produces highly valued nuts.

**Figure 3.**
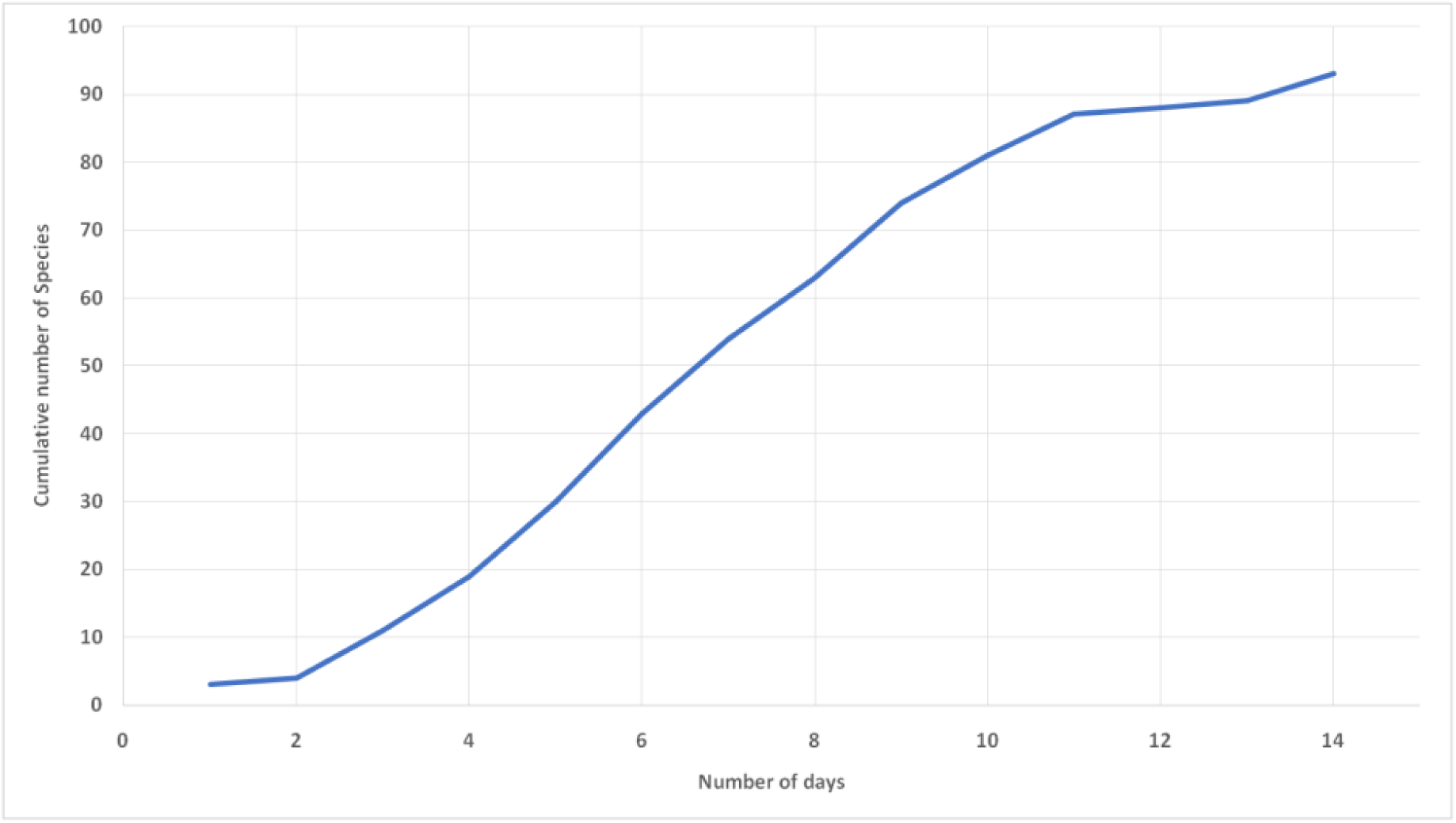
Species accumulation curve of wild edible plants mentioned during 14 days of walking in the forest with 20 informants around Le Bosquet and Kungu, southeast Cameroon, 2019.

### Species characteristics

The 91 wild edible plant species belonged to 43 different plant families, of which the best represented were Dioscoreaceae (ca. 9 species of wild yams), Irvingiaceae (8 spp.), Anacardiaceae (5 spp., including 4 species of *Trichoscypha* fruits) and Zingiberaceae (5 spp. of *Aframomum*).

Most wild edible plant species collected by the Baka naturally occur in primary forest (Figure 4). We encountered very little primary forest that was untouched by loggers: the only patch of forest that did not show signs of commercial timber harvesting was dominated by *Gilberiodendron dewevrei* (De Wild.) J. Leonard, located at ca. two hours walking distance from Le Bosquet. The selectively logged primary forest, however, contained the majority of the fruit and seed producing primary trees and lianas sought after by the Baka.

**Figure 4.**
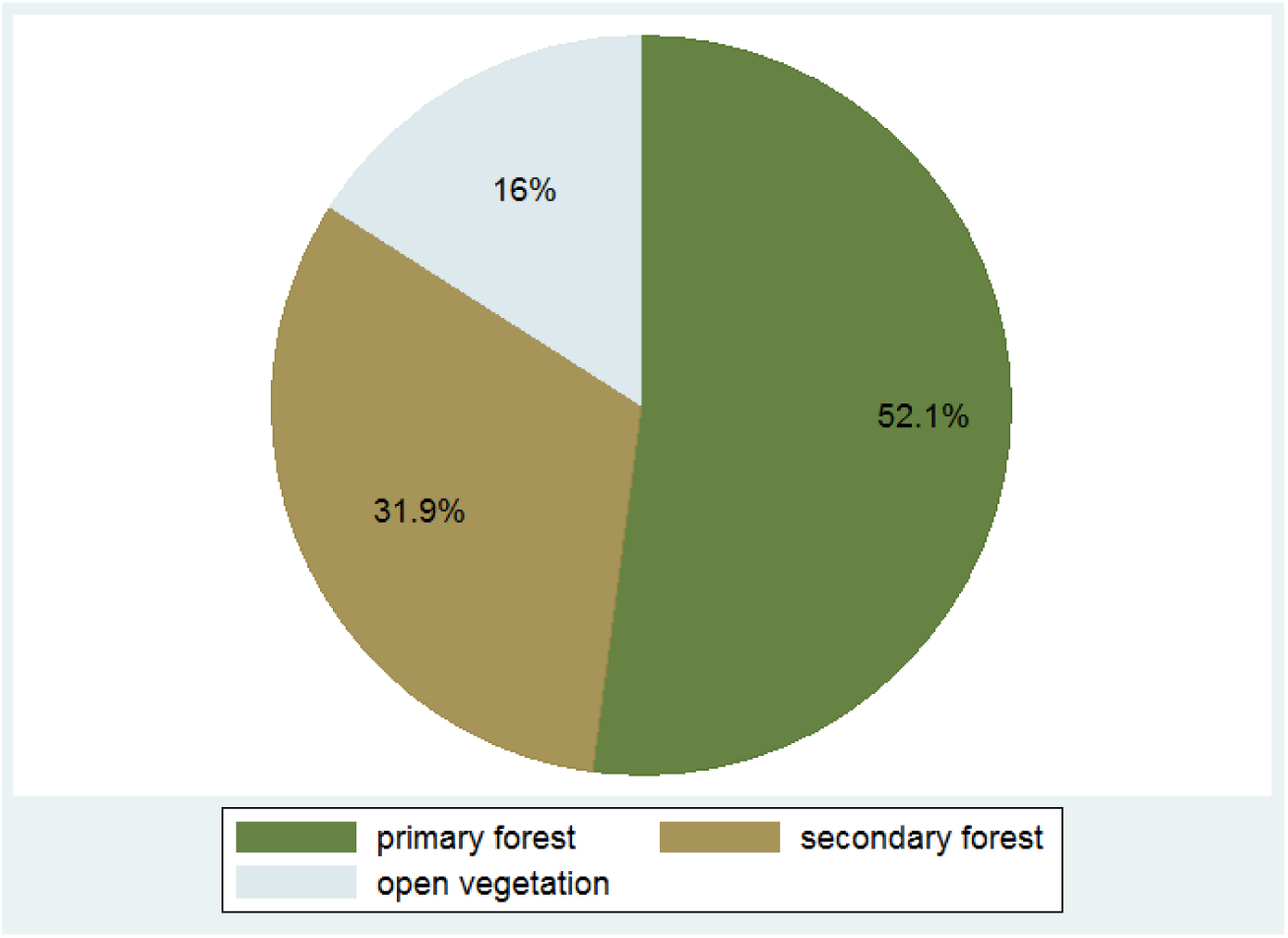
Distribution of the natural habitats of the wild edible plants (94 taxa) that were reported through the walk-in-the-woods method.

Most WEP-producing species were trees, followed by climbers; including both woody lianas and non-woody vines (Figure 5). Fruits (37%) and seeds (27%) were the most frequently mentioned edible plant parts, followed by leaves (19%), tubers (12%), bark (5%) and exudate (1%).

**Figure 5.**
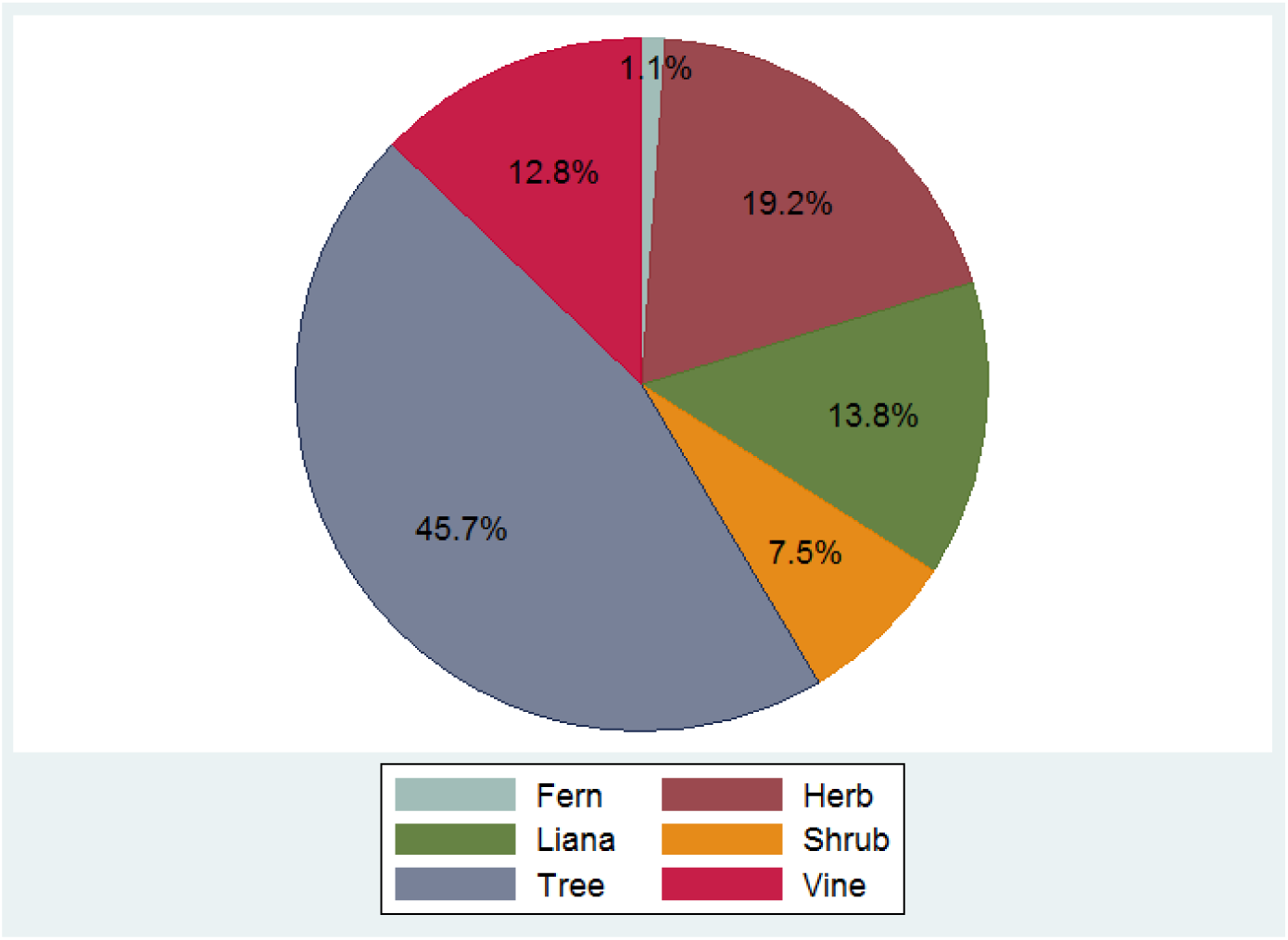
Distribution of the life forms of the species collected (94 species). Data collected through walk-in-the-woods method.

### Differences in WEP consumption data according to methodology

Of the 82 WEP species for which we had information on last consumption, 26 species were eaten within the last month by at least one of our 20 informants participating to the walk-in-the-wood expeditions (Figure 6), while 36 species were eaten within the last 12 months, 11 species between one and two years ago, eight species more than two years ago and one species was never eaten by any of our 20 informants.

**Figure 6.**
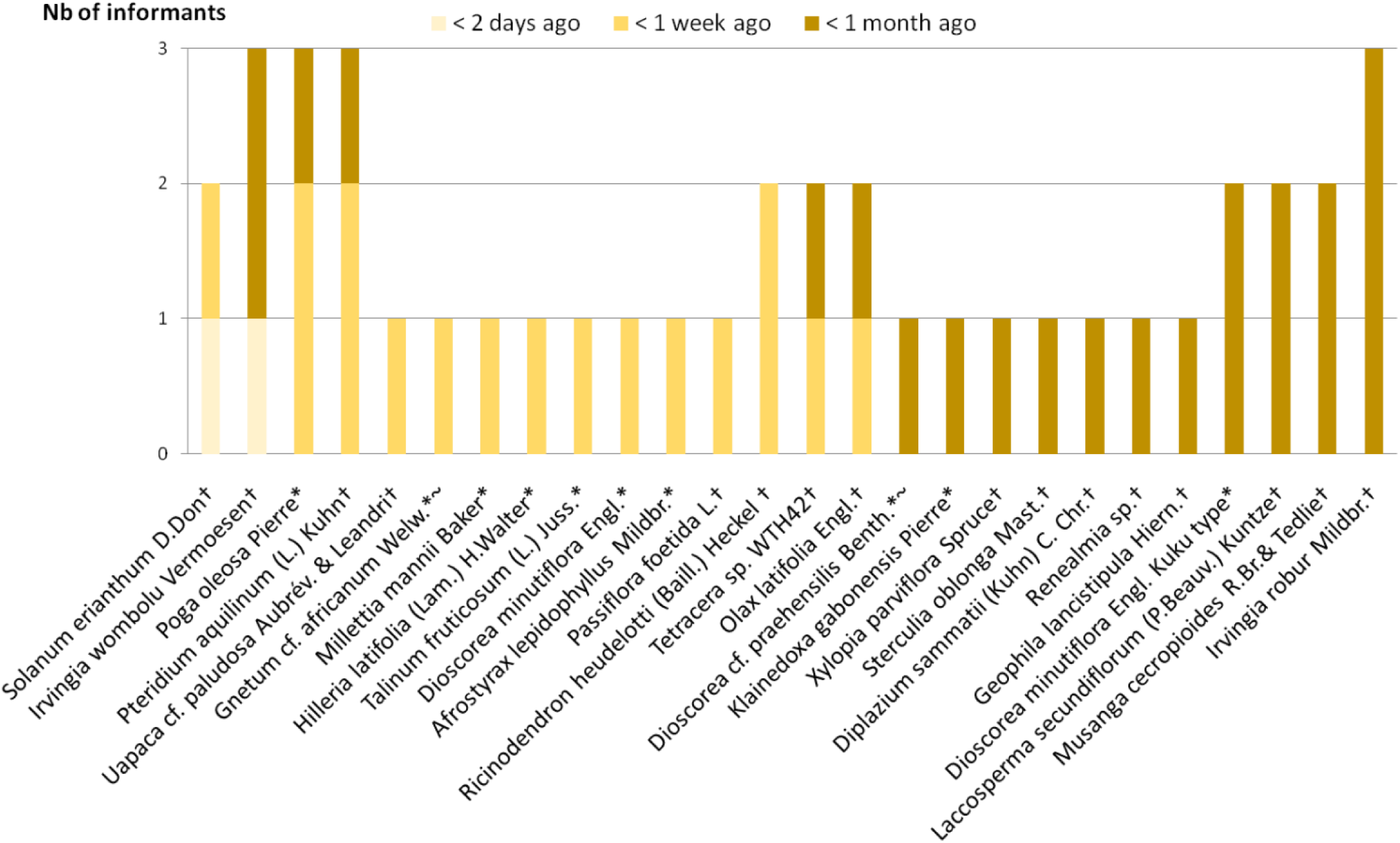
Wild edible species that our 20 informants consumed within the past month. * Species also reported through freelisting; ~ species also reported through dietary recalls, † species not reported either in freelisting or dietary recalls.

Only ten of the 26 species mentioned as recently consumed during the walk-in-the-woods method were reported during the freelisting and only three of these species also emerged through the dietary recalls, although the number of people interviewed during the last two methods was substantially higher. In other words, 23 recently consumed species would not have been identified with dietary recalls only, and 16 would have been missed if only the freelisting and dietary recalls would have been performed. These 16 species were two edible ferns (*Pteridium aquilinum* (L.) Kuhn and *Diplazium sammatii* (Kuhn) C. Chr.); three spices (*Xylopia parviflora* Spruce, *Olax latifolia* Engl., and *Ricinodendron heudelotti* (Baill.) Heckel); four fruits (*Passiflora foetida* L., *Solanum erianthum* D. Don, *Musanga cecropioides* R.Br. ex Tedlie and *Uapaca* cf. *paludosa* Aubrév. & Leandri); one inner stem (*Laccosperma secundiflorum* (P.Beauv.) Kuntze); three seeds (*Sterculia oblonga* Mast., *Irvingia robur* Mildbr. and *I. wombulu* Vermoesen); one tuber (*Renealmia* sp. WTH64); one drinkable water from the stem (*Tetracera* sp. WTH42) and one edible leaf which was eaten as a luck charm (*Geophila lancistipula* Hiern) (See Figure 6).

The most frequently mentioned WEP during the dietary recall was *Gnetum africanum* Welw., of which the leaves had the highest consumption during the major dry season, followed by several species of wild yams (*Dioscorea* spp.) and bush mango kernels (*Irvingia* spp.). Typically, the WEP most recently consumed by the participants during the forest surveys was the weedy shrub *Solanum erianthum*, of which the bitter fruits were boiled with wild garlic bark (*Afrostyrax lepidophyllus*) and (cultivated) hot pepper (*Capsicum frutescens*) and taken as a hot drink to wake up in the morning.

### Conflicts between wild fruit collection and commercial logging

During our forest walks, we identified six WEP of which the wood was observed as logged or said to be logged by the Baka (see Table 1). Three of these species were considered as vulnerable by the IUCN but none appeared on the CITES Appendixes I or II (Table 1). If only using ex situ interviews, we would have missed four species that are eaten by the Baka and also logged. Indeed, only two of these WEP-producing commercial hardwoods were mentioned in the freelisting interviews (*Baillonella toxisperma* and *Chrysophyllum lacourtianum*), while only one of them was recorded through the dietary recalls (*B. toxisperma*).

**Table 1.**
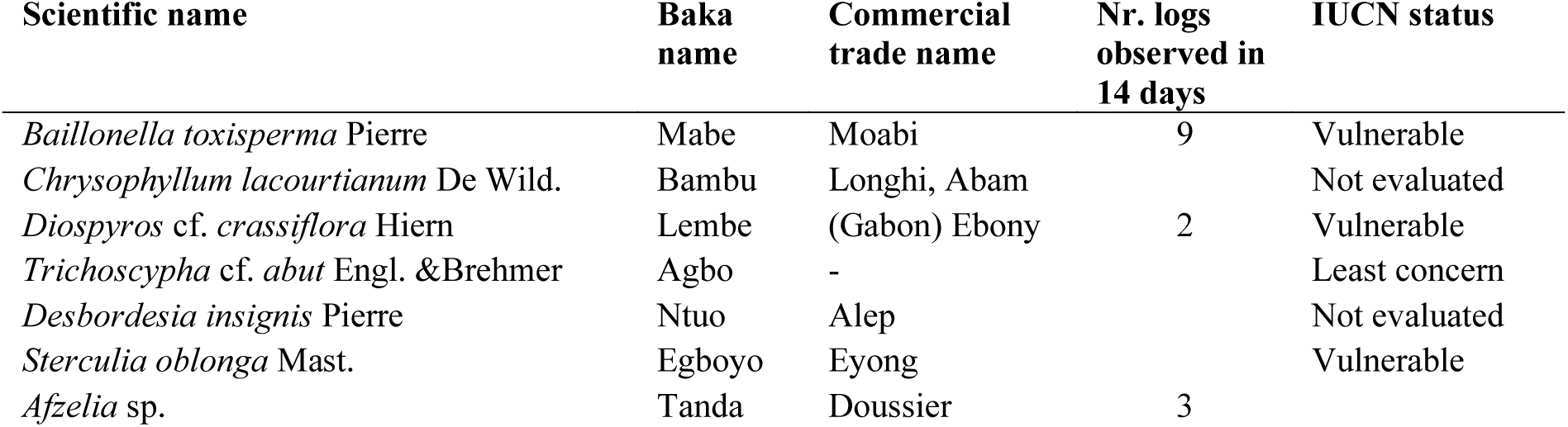
Commercial hardwood tree species producing edible fruits and/or seeds consumed by the Baka, trade names and current conservation status.

According to our informants, the moabi tree (*B. toxisperma*), highly valued by the Baka for their fresh fruits and seed oil, was on the most sought after by the logging companies operating in the Baka territory. Our informants mentioned that only trees exceeding 1 meter in diameter were felled, so several smaller individuals were still present. Other species that we observed as felled trunks, either along forest trails or on trucks in the 14 days were *Entandrophragma cylindricum* (Sprague) Sprague (four trunks), *Pterocarpus soyauxii* Taub.(five), *Piptadeniastrum africanum* (Hook.f.) Brenan (four), *Cylicodiscus gabunensis* Harms (one), *Rodognaphalon brevicuspe* (two) and *Triplochiton scleroxylon* K.Schum. (six). Although these are inedible species, they have several uses in (ritual) medicine, and *E. cylindricum* commonly hosts edible caterpillars, an important food for the Baka. During the forest walks, we also observed several (smaller) trees cut down by the Baka themselves, mostly to obtain fresh leaves of *Gnetum* cf. *africanum* lianas, to harvest honey, and once to collect the bitter bark of *Garcinia kola* Heckel., which is added to *Raphia* palm wine as a flavoring agent.

### Commercial Non-Timber Forest Products revealed through the different methods

During the walk-in-the-wood surveys, the Baka pointed out 24 different WEP species that they sold to middlemen, mostly in the form of fruits, seeds, or the oil from seeds (Table 2). During the earlier conducted interviews on the general income from sale over 14 days, only six different taxa were reported to have been sold.

**Table 2.**
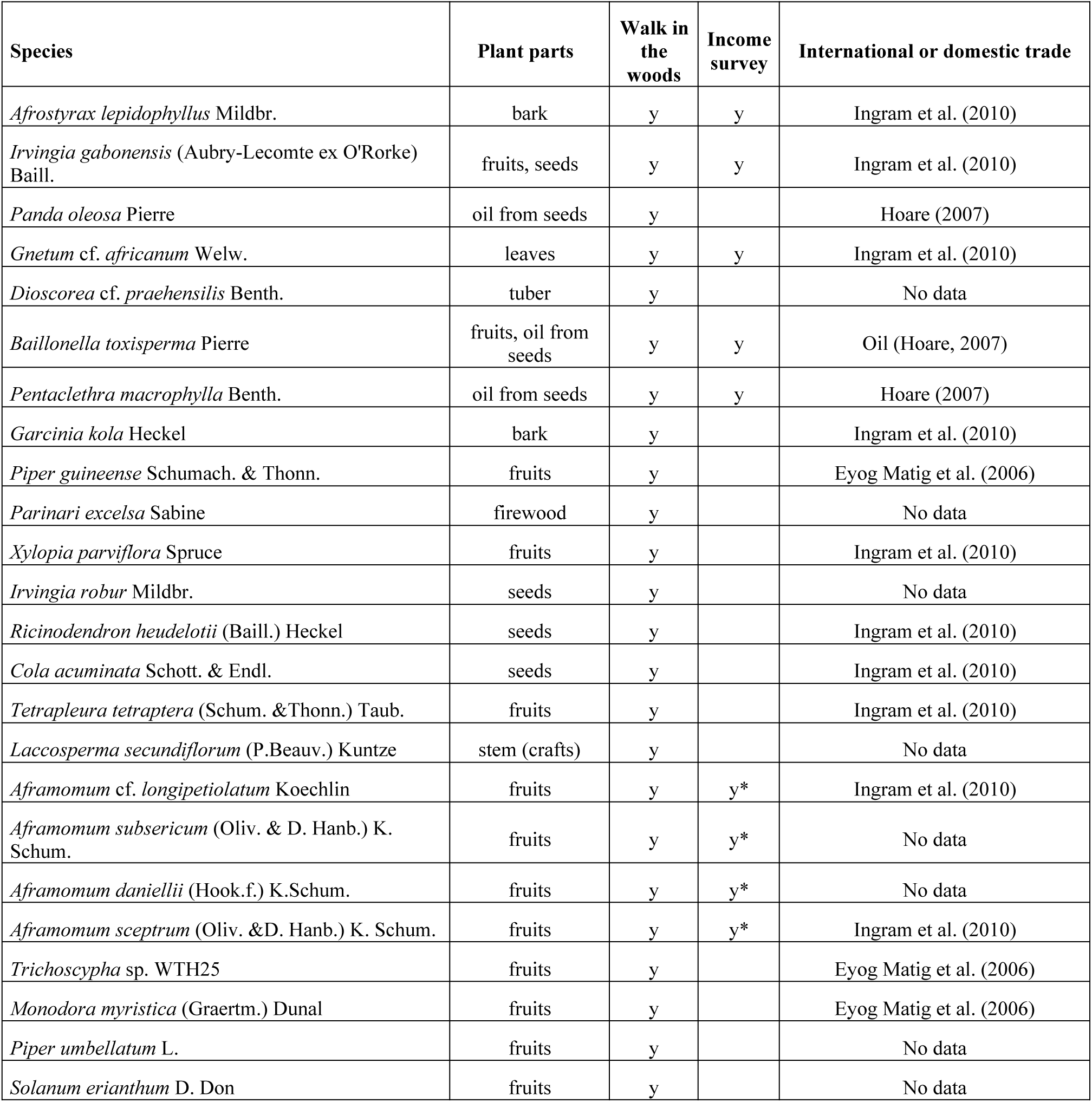
Data on wild food plant products sold by the Baka, retrieved through different methods. *Only “tondo”, the general Baka term for *Aframomum* sp. was reported in the income survey.

The most frequently sold species was *Gnetum* cf. *africanum*, followed by the seeds and oil of *Irvingia gabonensis, B. toxisperma, Pentaclethra macrophylla, Afrostyrax lepidophyllus* bark and the fruits of several unspecified *Aframomum* species. In the same line, of these 24 commercial NTFP, only six species were reported in the dietary recalls, and eight during the freelisting. Of the 24 commercial species that appeared during the forest surveys, 13 are commonly sold on the international market (Eyog Matig et al., 2006; Hoare, 2007; Ingram & Schure, 2010). The importance of these NTFP for the Baka livelihood, either for home consumption or (inter-) national trade would have been missed when our research methods had been limited to *ex-situ* interviews, the 14 day income recalls. The extraction of large moabi trees (*B. toxisperma*) by commercial timber companies must affect the amount of fruits and seeds that remain available for the Baka’s subsistence and cash income.

## Discussion

Although our research was performed among a relatively small population, our results show that different methods resulted in substantial differences in the collected data. *Ex-situ* interviews did not capture the full diversity of wild edible plants known, used and sold by the Baka. This may be partly due to the fact that not every participant understood the concept of “wild edible plant”, as this does not have a literal translation in Baka language, and the phrasing “food from the forest excluding game, honey and mushrooms” had to be used (Gallois et al., 2020). Wild food plants play an important role in Baka livelihood (Bahuchet, 1992; Dounias, 1993) and knowledge related to edible plants is acquired early during childhood (Gallois et al., 2017). Therefore, it seems unlikely that the Baka adult informants who did not report any WEP during the freelisting did not know any; they probably did not understand the domain.

During the walk-in-the-woods method, the researcher can directly exclude items pointed out by informants that fall outside the domain ‘wild edible plant’, such as fungi, animal products and cultivated plants, although the latter category can be challenging due to the presence of wild species under various degrees and types of human management and intervention through to domestication (Bharucha & Pretty, 2010). The advantage of assessing plant knowledge within the ecological context is that many species are encountered that do not pop-up quickly in people’s minds during a (shorter) interview outside the forest. When walking through the natural environment where edible plants occur, it is easier to remember them because of the amount of visual references to this knowledge at that moment (Miranda et al., 2007).

Our walk-in-the-woods method resulted in a higher number of wild edible species (even with a small sample size of only 20 informants) and elicited 12 recently consumed species that did not appear through the dietary recalls. We speculate that these were plants that were easily forgotten (e.g., spices, condiments, small fruits), species that people might feel ashamed of eating (e.g., ferns, weedy plants), or items that were previously missed due to misinterpretation of the term “wild edible plant” (e.g., drinking water from lianas, edible latex, ritual food plants) during the free listings. Additionally, our botanical inventory through forest walks revealed that some general terms for local taxa mentioned during interviews actually included several species. In the case of the Baka, the local name ‘tondo’ may refer to three different species of *Aframomum*, the term ‘bokoko’ to two species of *Klainedoxa*, and ‘payo’ to different species of *Irvingia* (see also Gallois et al., under review).The walk-in-the-woods method, however, is laborious to perform and requires additional botanical collection, as more rare species will be encountered that are hard to identify, for which the help of taxonomic specialists and support from herbaria is needed. The number of informants that can be taken into the field is also limited, while freelisting exercises can be organized in a short time among larger numbers of people (Paniagua Zambrana et al., 2018). On the other hand, *ex-situ* interview methods (including freelisting and dietary recalls) are known to assess the most salient useful plants among a large group of people in a relatively short time, as these techniques are limited by their spatio-temporal context (De Sousa et al., 2016; Paniagua Zambrana et al., 2018).

The implications of studies based solely on *ex-situ* interviews can be serious, as they lead to an underestimation of wild edible plants known, consumed, and commercially exploited, either by the local population or by outsiders. Results of such studies may not be representative for the situation on the ground, as trade in NTFP or conflicts between wild fruit collection and logging of fruit-producing trees may remain invisible. Moreover, the assessment of the contribution of wild plants to local diet and nutrition may be inaccurate. Several studies based on dietary recalls have concluded that WEP do not play an important role in local diets. For instance, Termote et al. (2012:8) were “confident to provide a fair representation of the dietary contribution of WEP on a population level in our sample”; even though their botanical collection was limited to finding specimens to match the local names mentioned during their dietary recalls and freelisting interviews. In Brazil, Do Nasciamento et al. (2013: 337) stated after their freelisting and dietary recall surveys that “The low consumption of wild species [….] is notable, which suggests that, in practice, these foods contribute little to contemporary dietary enrichment”. Such data could be misused by policy makers, who may conclude that rural communities do not need the forest that much as previously thought which seriously underestimates their use and dependency of forest resources.

Considering the importance of wild plants for food security and for providing nutrients that are not present in other foods (Ong & Kim, 2017), and the fact that children are major consumers of wild fruits but hardly recruited as interviewees (Guinand & Lemessa, 2000; Setalaphruk & Price, 2007), it is crucial to draw the most accurate overview of the diversity of wild food items used by local people. The various direct and indirect effects of logging and trade in NTFP may impact not only human food resources, but the entire ecosystem. Many of the oily seed producing trees in Central Africa are ecological keystones species that are crucial for the survival of local wildlife (Beaune et al., 2013), on which forest-dwelling groups such as the Baka rely on for meat.

## Conclusion

As expected, our walk-in-the-woods method resulted in a much higher number of wild edible plant species than the dietary recalls and freelisting methods, but species reported as most frequently consumed differed between the three methods. Our hypothesis that the list of plants generated by freelisting and recalls methods (either dietary or income) underestimated (conflicting) uses of edible plants proved to be correct. Our mixed methods approach shows the importance of cross-referencing data, not only between different types of interviews, as recommended by Paniagua-Zambrana et al. (2018), but also between interviews and direct observation during forest trips, for a better assessment of the diversity, consumption frequency and conflictive uses of wild edible plants. We therefore recommend that wild plant knowledge and use should be assessed through an “open” walk-in-the-woods method, in which informants are encouraged to mention any useful plant they know or randomly encounter, after which they are asked when they last used it. Employing the walk-in-the-woods technique merely to supply specimens for previously composed lists of useful plants from literature or interviews limits the capacity of this powerful technique to assess wild plant knowledge and use. Freelisting and dietary recalls can be used afterwards to supplement the walk-in-the-woods results with additional quantitative data, but they should not limit it, especially in the case when biased conclusions may have large implications for people’s future wellbeing.

## Acknowledgments

This research has received funding from the European Research Council under the European Union’s Horizon 2020 research and innovation program; grant agreement number STG–677576 (“HARVEST”). The botanical fieldwork was funded by Naturalis Biodiversity Center, the TreubMaatschappij and the Alberta Mennega Stichting. We would like to thank our colleagues, especially field assistants Appolinaire Ambassa and Ernest Isidore Simpoh and driver Alain Hyppolite Fezeu for their help with the fieldwork. We deeply thank Professor Bonaventure Sonké for his support in Cameroon. David Harris (Edinburgh herbarium), Jan Wieringa, Paul Maasand Carel Jongkind (Naturalis) and Marc Sosef (Meise Botanical Gardens), helped us to identify our specimens. Finally, our greatest gratitude goes to all Baka children, women and men with whom we have lived and worked. Thank you for your trust, hospitality and generous hearts.

## Author Contributions Statement

SG, AGH and TVA designed the study, SG, WTH and TVA collected the data. WTH conducted the first analysis and wrote the first manuscript for his Msc thesis. All the authors then elaborated the current manuscript, from data analysis to writing the final version.

## Contribution to the Field Statement

Many small-scale human societies rely on their access to natural resources for their daily diet. Due to globalization and forest degradation, many of them are undergoing a nutritional transition, in which there is an increase of fat- and sugar-rich processed foods, at the expense of wild plants. To assess the wild edible plants known and consumed by local people, several studies have used *ex-situ* interview methods, such as freelisting and 24h dietary diversity recalls. In our study, we compared four different methods (freelisting, dietary recalls, income recalls and the walk-in-the-wood method) to explore how they differed in results with regard to the diversity, consumption frequency and conflictive uses of wild edible plants. Working with Baka forager-horticulturalists in southeastern Cameroon, we showed that dietary recalls and freelisting strongly underestimate people’s knowledge and consumption of wild plants. These insights raise questions on what can be interpreted from *ex-situ* interviews, as well as the possible scientific and political consequences of misinterpreting data on the wild food resources for forest-dwelling people.

## Footnotes

1 (http://plantsoftheworldonline.org/).

2 https://www.cites.org/eng/app/appendices.php

3 https://www.iucnredlist.org

4 https://www.itto.int/

